# Prenatal thyroid hormone exposure increases growth but not oxidative stress in a wild passerine species

**DOI:** 10.1101/578047

**Authors:** Tom Sarraude, Bin-Yan Hsu, Ton G.G. Groothuis, Suvi Ruuskanen

## Abstract

Hormones transferred from mothers to their offspring are thought to be a maternal tool for mothers to prepare their progeny for expected environmental conditions, thus increasing fitness. Thyroid hormones (THs) are crucial across vertebrates for embryonic and post-natal development and metabolism. Nevertheless, the studies that investigated the consequences of maternal hormones have mostly focused on steroid hormones and ignored maternally-derived thyroid hormones. In this study, we experimentally elevated yolk thyroid hormones in a wild population of a migratory passerine, the European Pied flycatcher *Ficedula hypoleuca*. We injected eggs with a mixture of T_4_ and T_3_ within the natural range of the species to assess its effects on hatching success, nestling survival, growth and oxidative status (antioxidant enzyme activity, lipid peroxidation and oxidative balance). We found no effects of yolk THs on egg hatchability or nestling survival. Yolk THs increased nestling growth during the second week post hatching, but this potentially beneficial effect did not incur any costs in terms of oxidative stress. The results should stimulate more research on thyroid hormone mediated maternal effects, further studies into the underlying mechanistic pathways for these effects and how they translate into adulthood and fitness.

**Summary statement:** Thyroid hormones have been overlooked in the context of hormone-mediated maternal effects. We found that yolk thyroid hormones in a wild bird species increase growth without incurring oxidative stress.

## Introduction

Maternal effects are all the non-genetic influences of a mother on her offspring and receive increasing attention in evolutionary and behavioural ecology. Via maternal effects, mothers may influence the fitness of their progeny by adapting their phenotype to expected environmental conditions (“adaptive maternal effects” in Marshall and Uller, 2007; Mousseau and Fox, 1998). Maternal effects are observed in plants, invertebrates and vertebrates, and can have many possible mediators (Danchin et al., 2011; Kuijper and Johnstone, 2018). One intriguing pathway is via the hormones transmitted from the mother to her progeny. These hormone-mediated maternal effects have been found to profoundly influence offspring phenotype in many different taxa (e.g., in mammals, Dantzer et al., 2013; birds, von Engelhardt and Groothuis, 2011; reptiles, Uller et al., 2007 and invertebrates, Schwander et al., 2008). Most studies in the field of hormone-mediated maternal effects have focussed on steroid hormones, especially cortisol, corticosterone and androgens (von Engelhardt and Groothuis, 2011). However, mothers transfer other hormones to their embryo (Williams and Groothuis, 2015), including thyroid hormones (Ruuskanen and Hsu, 2018) that deserve much more attention.

Thyroid hormones (THs) are metabolic hormones produced by the thyroid gland and are present in two main forms: the prohormone thyroxine (T_4_) and the biologically active form triiodothyronine (T_3_). Thyroid hormones have pleiotropic effects that serve several biologically important functions across vertebrates (Ruuskanen and Hsu, 2018). In early-life, they participate in the maturation of multiple tissues (e.g., birds, McNabb and Darras, 2015; mammals, Pascual and Aranda, 2013), and interact with growth hormones to increase growth (McNabb and Darras, 2015). THs also regulate metabolism, and, during adult life, influence reproductive functions (e.g., birds, McNabb and Darras, 2015; mammals, Norris and Carr, 2013). Because of their stimulating effect on metabolism, THs contribute to the production of reactive oxygen species (ROS) (Villanueva et al., 2013). Oxidative stress occurs when the ROS production exceeds the capacity of antioxidant defences (Monaghan et al., 2009). It results in oxidative damages on, for example, DNA, lipids and proteins (Monaghan et al., 2009). In mammals, for instance, THs augment the degree of unsaturation of the fatty acids in the mitochondrial membranes (Villanueva et al., 2013), which will make the membranes more prone to lipid peroxidation. Lipid peroxidation can ultimately alter mitochondrial function (Hulbert et al., 2007; Monaghan et al., 2009). In contrast, THs can activate antioxidant systems in the cell and can act themselves, at least *in vitro*, as free radical scavengers (Oziol et al., 2001). Despite such dual effects towards opposite directions, a general trend emerges from the literature with hyperthyroid tissues exhibiting higher oxidative damage (reviewed in Villanueva et al., 2013).

Studies on the effect of maternal thyroid hormones on offspring development in wild animals are scarce. In humans and rats hypothyroid condition of the mother impairs brain development and cognition in her children (Moog et al., 2017). A potential problem here is that in mammalian species, maternal thyroid variation or manipulation inevitably influences other aspects of maternal physiology, which confounds the direct effects on the offspring. Oviparous species, such as birds, are therefore suitable models for studying the role of maternal hormones on the progeny because embryos develop in eggs outside the mother’s body and maternally-derived hormones are deposited in egg yolks (Prati et al., 1992; Schwabl, 1993). This allows the measurement and experimental manipulation of maternal hormone transfer to be independent of maternal physiology. Together with their relatively well-known ecology and evolution, birds have become the most extensively studied taxa in research on the function of maternal hormones (Groothuis and Schwabl, 2008; von Engelhardt and Groothuis, 2011).

Maternal thyroid hormones have long been detected in egg yolks of chicken (Hilfer and Searls, 1980; Prati et al., 1992) and Japanese quail (Wilson and McNabb, 1997), although these hormones have received very little attention (Ruuskanen and Hsu, 2018). To date, only three studies have investigated the effects of physiological variations in yolk THs on offspring development (great tits, *Parus major*, Ruuskanen et al., 2016; rock pigeons, *Columba livia*, Hsu et al., 2017; collared flycatchers, *Ficedula albicollis*, Hsu et al., 2019). These studies revealed potential biological relevance and fitness consequences but also some discrepancies on the role of yolk THs. For example, yolk THs improved hatching success in rock pigeons (Hsu et al., 2017) and in collared flycatchers (Hsu et al., 2019) but had no effect in great tits (Ruuskanen et al., 2016). Moreover, THs injection in great tits increased growth in males but decreased it in females (Ruuskanen et al., 2016). Conversely, yolk THs decreased growth during the second half of the nestling phase in rock pigeons (Hsu et al., 2017), whereas they increased early growth in collared flycatchers (Hsu et al., 2019). Finally, great tits showed no response to elevated yolk THs in resting metabolic rate (RMR) (Ruuskanen et al., 2016), whereas a sex-specific effect on RMR was detected in rock pigeons (Hsu et al., 2017). These studies suggest that yolk THs can exert costs and benefits on the offspring in a species- or context-specific manner, or that these differences are due to different dosages of the hormone. Therefore, further studies on other species are needed to complement these findings. The study on collared flycatchers is the only one so far that investigated the association between yolk THs and oxidative stress in offspring (Hsu et al., 2019). This study surprisingly showed no adverse effect of yolk THs on oxidative damage or oxidative balance, despite the growth-enhancing effects (Hsu et al., 2019). This absence of influence on oxidative stress contradicts the general knowledge of THs, with hyperthyroid tissues exhibiting higher oxidative damage (Venditti and Meo, 2006; Villanueva et al., 2013), calling for additional studies to confirm or contradict these findings.

The aim of this study was to evaluate the short-term effects of maternal THs on the traits discussed above: hatching success, growth and nestling oxidative stress, using a closely related species to the above-mentioned collared flycatcher, the european pied flycatcher (*Ficedula hypoleuca*). We manipulated the concentrations of yolk THs in a wild population of this avian species, by injecting a combination of T_4_ and T_3_ in their eggs. In contrast to some earlier studies we ensured that the treatment was within the physiological range of the species.

## Material and Methods

### Study site and study species

The experiment was conducted during the spring 2017 in Turku, South-West of Finland (60°26’N, 22°10′E). The study species is the Pied flycatcher, a small (ca. 15 g) migratory passerine that breeds in Finland from May to July. Pied flycatchers are secondary cavity nesters that also breed in artificial nest boxes. At this latitude, females generally lay a single clutch of 5 to 8 eggs.

### Nest monitoring and experimental design

Yolk thyroid hormone concentrations were elevated via injections into unincubated eggs using a between-clutch design. In total, 29 clutches (170 eggs) received a thyroid hormone injection (hereafter TH-treatment), and 28 clutches (169 eggs) received a control injection (hereafter CO-treatment). In two nests, one in each treatment, none of the eggs hatched due to desertion before incubation. These two clutches were therefore removed from the analysis. The final sample size is 28 TH-nests (166 eggs) and 27 CO-nests (164 eggs).

Nest boxes were monitored twice a week during nest construction until egg laying. On the morning when the fifth or sixth egg was laid, all eggs were temporarily removed from the nest for injection, replaced by dummy eggs and returned after injection. Nests were then visited every following morning to inject freshly laid eggs until clutch completion, marked by the absence of freshly laid eggs and females incubating their eggs.

The clutches were randomly assigned to one of the treatments. In addition, treatments were alternated across clutches to balance the order of treatments within a day. Similarly, we also balanced the treatments across the laying period. There was no difference in the average (± SD) laying date (TH = 27.00 ± 2.64 vs. CO = 27.19 ± 2.65, 1 = 1^st^ of May, Wilcoxon unpaired test, W = 402.5, p = 0.68), nor in the average (± SD) clutch size (TH = 5.93 ± 0.81 eggs vs. CO = 6.07 ± 0.78 eggs, Wilcoxon unpaired test, W = 439.5, p = 0.26).

### Preparation of the solution and injection procedure

The thyroid hormone solution (TH solution) was composed of a mix of T_4_ (L-thyroxine, ≥ 98% HPCL, CAS number 51-48-9, Sigma-Aldrich) and T_3_ (3,3’,5-triiodo-L-thyronine, >95% HPCL, CAS number 6893-02-3, Sigma-Aldrich), first dissolved in 0.1M NaOH and then diluted in 0.9% NaCl. The concentration of each hormone was based on hormone measurements in 15 pied flycatcher eggs, from 15 clutches, collected during the spring 2016 in Turku in which the average hormone contents were the following: T_4_ = 2.307 ng/yolk (SD = 0.654) and T_3_ = 0.740 ng/yolk (SD = 0.238). The injection of thyroid hormones resulted in an average increase of two standard deviations, a standard and recommended procedure for hormone manipulation within the natural range (Hsu et al., 2017; Podmokła et al., 2018; Ruuskanen et al., 2016). The control solution (CO) was a saline solution (0.9% NaCl).

Before the injection, the shell was disinfected with a cotton pad dipped in 70% alcohol. The injection procedure consisted of four steps. First, a disposable and sterile 25G needle (BD Microlance ™) was used to pierce the shell. To locate the yolk, the egg was lit by a small torch from underneath. Second, the injection of 5µl was performed with a Hamilton^®^ syringe (25 µl, Hamilton Company) directly into the yolk. Third, the hole in the shell was sealed with a veterinary tissue adhesive (3M Vetbond ™) and the eggs were marked with a permanent marker. Finally, all eggs from the same clutch were returned to the nest at the same time, and the dummy eggs removed.

### Nestling growth monitoring and blood sampling

Nests were checked daily for hatching two days before the expected hatching date. The date of hatching for a particular nest was recorded as the day the first hatchlings were observed (day 0). Two days after hatching, nestlings were coded by clipping feathers to identify them individually. Nestlings were ringed at day 7 after hatching. Body mass (0.01 g) was recorded at day 2, 7 and 12 after hatching. Tarsus (0.1 mm) and wing length (1 mm) were recorded at day 7 and 12. At day 12 blood samples were also collected (ca. 40 µl) from the brachial vein in heparinized capillaries and directly frozen in liquid nitrogen for analyses of oxidative stress biomarker and molecular sexing. Samples were stored at −80°C until analyses. Finally, fledging was monitored from day 14 after hatching. Fledging date was recorded when all the nestlings had fledged from the nest.

### Sexing method

DNA extraction procedure from the blood cells followed Aljanabi and Martinez (1997). Primer R1 and F2 were used in PCR to amplify ATP5A1 gene located on avian sex chromosomes (Bantock et al., 2008). Overall, the method followed that described by Ruuskanen and Laaksonen (2010), with minor changes. The PCR condition for each sample was as following: 5 µl QIAGEN multiplex PCR kit + 0.1 µl of each primer (20 µM) + 1.8 µl H2O + 3 µl DNA, yielding 10µl for the final PCR volume. The initial denaturation was at 95 °C for 15 min, followed by 35 cycles of 95°C for 30 s, 55 °C for 90 s, and 72 °C for 60 s. The samples were then held at 72 °C for 10 min and 20 °C for 5 mins. PCR products were analysed with 3% agarose gel under 100 V for 90 min.

### Oxidative stress analysis methods

Samples from two individuals per clutch were analysed due to practical limitations. Whenever possible, one male and one female were chosen of approximately the same body mass. If samples from both sexes were not available for a clutch, then two individuals of the same sex were selected. In total, 103 nestlings were included in the analysis (TH, N = 27 nests and 50 nestlings; CO, N = 27 nests and 53 nestlings).

Three biomarkers of oxidative status were measured: the activity of the antioxidant enzyme glutathione S-transferases (GSTs), the ratio of reduced and oxidised glutathione (GSH:GSSG ratio) and lipid peroxidation (using malonaldehyde, MDA, as a proxy). GST enzymes catalyse the conjugation of toxic metabolites to glutathione (Halliwell and Gutteridge, 2007; Sheenan et al., 2001). In normal cells, GST activity is expected to be lower than in damaged cells (Rainio et al., 2013). The GSH:GSSG ratio represents the overall oxidative state of cells, and a low ratio reveals oxidative stress (e.g., Halliwell and Gutteridge 2007, Rainio et al. 2013, 2015). Lipid peroxidation is commonly measured with the thiobarbituric acid test (TBARS, (Alonso-Alvarez et al., 2008; Halliwell and Gutteridge, 2007). This test relies on the ability of polyunsaturated fatty acids (PUFAs) contained in cell membranes to readily react with oxygen radicals by donating a hydrogen atom. The fatty acid radical is unstable, and a chain of reactions occurs. Malonaldehyde (MDA) is an end product of this reaction (Marnett, 1999) and thus used as a measure of lipid peroxidation.

Whole blood was first diluted in 0.9% NaCl to achieve protein concentrations ranging 4–13 mg/ml. Overall protein concentration (mg/ml) was measured using BCA protein assay (Thermo Scientific) with a BSA standard (bovine serum albumin, Sigma). The methodology for measuring each GST and GSH:GSSG ratio followed Rainio et al. (2015). In brief, GST-assay (Sigma CS0410) was adjusted from 96-to 384-well plate. 2 μl of each sample were used in triplicates and our own reagents; Dulbecco’s phosphate buffered saline-buffer, 200 mM GSH (Sigma G4251), 100 mM CDNB (Sigma C6396) in EtOH. The GSH:GSSG ratio was measured with the ThioStar® glutathione detection reagent kit (K006-F1D Arbor Assays, USA) according to the kit instructions. The marker of lipid peroxidation, MDA, was analysed using a 384-plate modification of TBARS-assay described by Espin et al. (2017). In brief, 50 µl of samples diluted in 0.9% NaCl were mixed with 100 µl of TBARS-BHT reagent (15% Trichloroacetic acid, TCA; 0.375% 2-Thiobarbituric acid, TBA; 0.02% Butylated hydroxytoluene, BHT; and 0.25 N hydrochloric acid, HCl) and incubated in a waterbath at 90°C for 30 min. Samples were then cooled in ice-water for 10 min to stop the reaction, and centrifuged for 15 min at 6 ºC and 2100 g. Standard was prepared from MDA (Sigma). The samples were analysed in black 384-well plates, and fluorescence intensity (FI) was measured at an excitation/emission wavelength of 530/550 nm (EnSpire microplate spectrofluorometer) to obtain the highest sensitivity. All biomarkers enzyme activities were measured in triplicate (intra-assay coefficient of variability [CV] <10% in all cases).

### Statistical analysis

Data were analysed with the software R version 3.5.1 (R core team, 2018). General and generalised linear mixed models (LMMs and GLMMs, respectively) were performed using the R package *lme4* (Bates et al., 2015). Covariates and interactions between predictors were removed from the models when non-significant (p-values > 0.05). P-values in LMMs were obtained by model comparison using Kenward-Roger approximation from the package *pbkrtest* (Halekoh and Højsgaard, 2014). The significance of the predictors in GLMMs was determined by parametric bootstrapping with 1,000 simulations using the package *pbkrtest*. Model residuals were checked for normality and homogeneity by visual inspection. Significant interactions were further analysed by post-hoc comparison with the package *phia* (de Rosario-Martinez, 2015). Estimated marginal means and standard errors (EMMs ± SE) were derived from models using the package *emmeans* (Lenth, 2019).

To analyse hatching success, a dummy code was given to each egg: 0 for unhatched egg and 1 for hatched egg. Fledging success was coded similarly: 0 for dead and 1 for fledged nestling. Two GLMMs were performed with a binomial error distribution (logit link) and treatment as the predictor.

Duration of the embryonic period and duration of the nestling phase were fitted in separate linear models with treatment as the fixed effect. Laying date and brood size were added as covariates for nestling phase duration as they both may influence nestling growth and thereby nestling phase duration (Williams, 2012).

Models to analyse the morphological variables included sex as a fixed factor to test for potential sex-dependent effects of THs, as found by Hsu et al. (2017) and Ruuskanen et al. (2016). Early body mass (i.e., at day 2 after hatching) was analysed separately from growth during the second week post-hatching (i.e., from day 7 to day 12) for two reasons. First, variation in the former represents more the influence of maternal THs on prenatal development, while the variation in the latter also reflects the influence during the postnatal stage when the yolk that contain the hormones is totally consumed. Second, including the three time points in a single model created a non-linear growth curve hampering proper statistical analyses. The model to analyse early body mass included laying date as a covariate. Models to analyse growth during the second week post-hatching (i.e., growth in body mass, tarsus and wing length between day 7 and 12) included age and treatment as fixed factors, brood size at day two as an additional covariate, and nestling identity as an additional random intercept. Two- and three-way interactions between treatment, age and sex were tested and presented when statistically significant. Full models are detailed in Table S1.

The models of oxidative stress biomarkers (i.e., GST activity, MDA concentration and GSH:GSSG ratio) were similar as above but included body mass at day 12, which was the day of blood sampling, as an additional covariate, because body mass is known to be associated with oxidative status (e.g., Rainio et al., 2015). Assay number was also added as a random intercept to account for inter-assay variation. Response variables were log-transformed to achieve normal distribution. Full models are detailed in Table S2.

### Ethical note

The study complied with Finnish regulation and was approved by the Finnish Animal Experiment Board (ESAVI/2389/04.10.07/2017) and by the Finnish Ministry of Environment (VARELY580/2017).

## Results

### Hatching and fledging success, duration of embryonic and nestling periods

Hatching success (TH = 75.3% vs. CO = 76.8%) and fledging success (TH = 92.2% vs. CO = 92.3%) were comparable between the two treatments (hatching success, GLMM, p = 0.682; fledging success, GLMM, p = 0.920). Duration of the embryonic period did not differ between the groups (mean ± SD TH = 13.8 ± 1.2 days vs. CO = 13.8 ± 1.3 days, t = −0.225, p = 0.823). Injection of yolk THs did not affect the duration of the nestling period either (mean ± SD TH = 15.5 ± 0.7 days vs. CO = 15.7 ± 0.9 days, t = −0.991, p = 0.327). Likewise, there was no effect of laying date on the duration of the nestling period (t = −0.371, p = 0.712). Brood size at day two after hatching reduced significantly the duration of the nestling period (E ± SE = −0.167 ± 0.083, t = −2.025, p = 0.049).

### Growth

We detected a significant interaction effect between treatment and age on nestling body mass between day 7 and 12 (Table 1). Post-hoc analyses revealed that TH-treated nestlings grew faster than control nestlings during the second week post-hatching (adjusted slopes ± SE = 0.43 ± 0.02 for CO nestlings, 0.49 ± 0.02 for TH nestlings, *Χ*^2^ = 4.85, Holm-adjusted p = 0.028, Figure). However, there was no significant difference in body mass between the treatments at day 7 (EMMs ± SE: CO = 11.83 ± 0.16 g vs TH = 11.70 ± 0.17 g, *Χ*^2^ = 0.36, Holm-adjusted p = 0.908) or day 12 (EMMs ± SE: CO = 13.96 ± 0.16 g vs TH = 14.14 ± 0.17 g, *Χ*^2^ = 0.56, Holm-adjusted p = 0.908), indicating that the interaction likely originates from small differences in the opposite directions at day 7 and day 12 between TH and control groups. On average, males had a larger body mass than females (EMMs ± SE: males = 12.99 ± 0.12 g, females = 12.82 ± 0.12 g, Table 1). For tarsus and wing lengths, however, no effects of yolk TH treatment were detected (Table 1). Experimental elevation of yolk thyroid hormones did not affect early postnatal body mass (EMMs ± SE: CO = 3.6 ± 0.1 g and TH = 3.5 ± 0.1 g). In addition, there were no sex differences in body mass at day 2 (Table 1).

**Table 1:** Results of linear mixed models of morphometric measures in response to yolk thyroid hormone elevation, in nestling pied Flycatchers (sample sizes: TH = 125; CO = 126). Measures were taken at day 2, 7 and 12 after hatching. Models presented here are the reduced models, after removing non-significant covariates and interactions. The predictors treatment and sex were retained in all models. Brood identity was included as a random intercept in all models, and nestling identity was included as a random intercept for the growth models between day 7 and 12. P-values and ddf were obtained using the Kenward-Roger approximation. Ndf = 1. Significant p-values are shown in bold.

|  |  |  |  |  |
| --- | --- | --- | --- | --- |
| **Body mass (g) at day 2** |  |  |  |  |
|  | Estimate | SE | F<sub>ddf</sub> | p |
| Treatment (TH) | -0.162 | 0.174 | 0.861<sub>51.8</sub> | 0.358 |
| Sex (Male) | 0.038 | 0.054 | 0.501<sub>181.9</sub> | 0.480 |
| **Body mass (g) from day 7 to 12** |  |  |  |  |
|  | Estimate | SE | F<sub>ddf</sub> | p |
| Treatment (TH) | -0.578 | 0.350 | 0.007<sub>51.6</sub> | 0.936 |
| Age (12 days) | 0.426 | 0.020 | 1016.8<sub>227.0</sub> | **<0.001** |
| Sex (Male) | 0.167 | 0.080 | 4.260<sub>185.1</sub> | **0.040** |
| Treatment (TH) × age | 0.063 | 0.028 | 4.849<sub>226.0</sub> | **0.029** |
| **Wing growth (mm) from day 7 to 12** |  |  |  |  |
|  | Estimate | SE | F<sub>ddf</sub> | p |
| Treatment (TH) | -0.186 | 0.569 | 0.107<sub>51.7</sub> | 0.745 |
| Age (12 days) | 4.704 | 0.025 | 36299<sub>227.0</sub> | **<0.001** |
| Sex (Male) | 0.199 | 0.188 | 1.115<sub>183.1</sub> | 0.293 |
| **Tarsus growth (mm) from day 7 to 12** |  |  |  |  |
|  | Estimate | SE | F<sub>ddf</sub> | p |
| Treatment (TH) | -0.074 | 0.108 | 0.466<sub>51.4</sub> | 0.498 |
| Age (12 days) | 0.287 | 0.006 | 2189.4<sub>227.0</sub> | **<0.001** |
| Sex (Male) | 0.007 | 0.048 | 0.023<sub>190.2</sub> | 0.881 |

**Figure 1:**
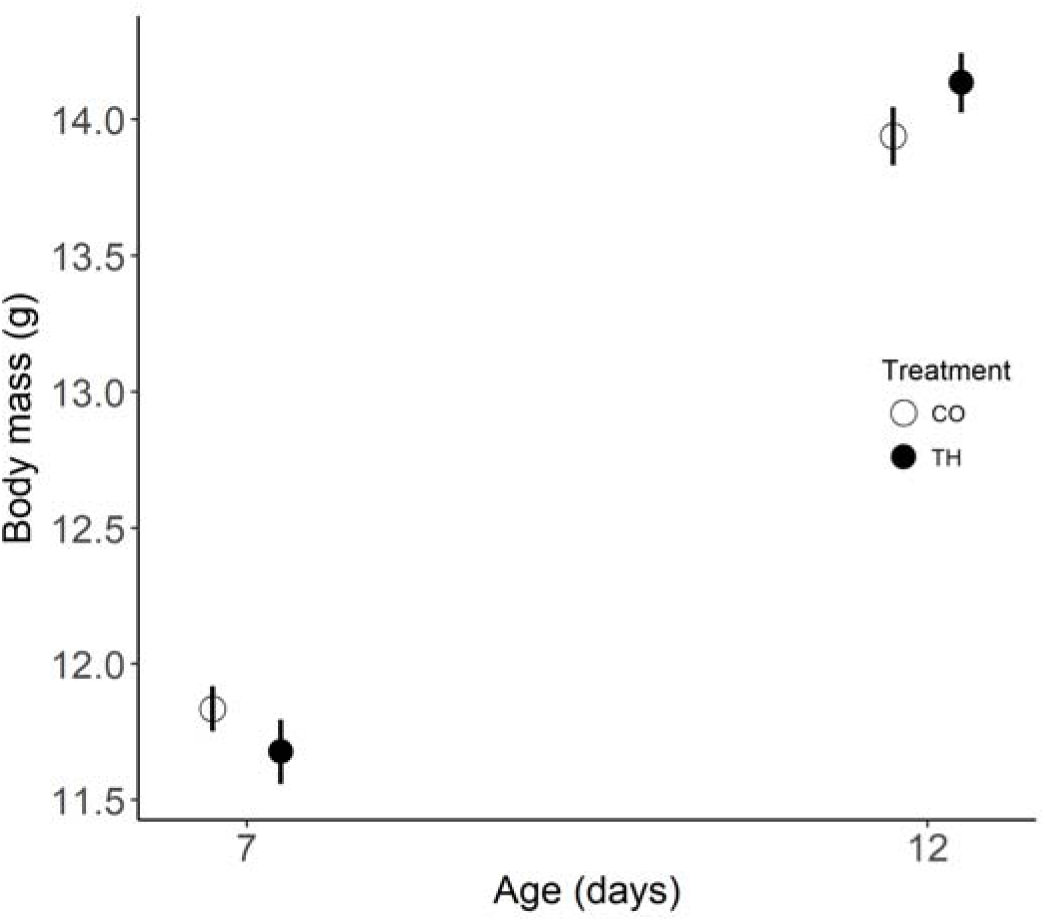
Body mass (mean ± SE) at day 7 and day 12 after hatching in TH-treated nestlings (N = 125) and controls (N = 126). Empty circle, CO = control injection; solid circle, TH = yolk thyroid hormone elevation. The interaction between the treatment and the age of the nestling was significant (Holm-adjusted p = 0.028).

### Oxidative stress and oxidative damage

Experimental elevation of yolk THs did not affect antioxidant enzyme activity (mean ± SE: CO and TH nestlings = 0.006 ± 0.0003 pmol GST/min/mg protein), oxidative damage on lipids (mean ± SE: CO = 0.051 ± 0.003 and TH = 0.053 ± 0.004 nmol MDA/mg protein) or oxidative status (mean ± SE: CO = 3.86 ± 0.47 and TH = 4.51 ± 0.73 GSH:GSSG ratio) (Table 2). In addition, neither sex of the nestlings nor their body mass had an influence on these oxidative stress biomarkers (Table 2). MDA concentration was negatively correlated with laying date (Table 2).

**Table 2:** Results of linear mixed models of oxidative stress biomarkers in response to yolk thyroid hormone elevation, in nestling pied Flycatchers at day 12 after hatching (TH = 50; CO = 53). Response variables were log-transformed to achieve normal distribution. Estimates and standard errors are on the log-scale. Models presented here are the reduced models, after removing non-significant covariates and interactions. The predictors treatment, sex and body mass at day 12 were retained in all models. Brood identity was included as a random intercept in all models. P-values and ddf were obtained using the Kenward-Roger approximation. Ndf = 1. Significant p-value is shown in bold.

| **GST activity (pmol/min/mg protein)** |  |  |  |  |
| --- | --- | --- | --- | --- |
|  | Estimate | SE | F<sub>ddf</sub> | p |
| Treatment (TH) | -0.073 | 0.058 | 1.591<sub>47.4</sub> | 0.213 |
| Sex (male) | -0.076 | 0.056 | 1.823<sub>59.4</sub> | 0.182 |
| Mass at Day 12 | 0.019 | 0.028 | 0.423<sub>52.7</sub> | 0.518 |
| **MDA concentration (nmol/mg protein)** |  |  |  |  |
|  | Estimate | SE | F<sub>ddf</sub> | p |
| Treatment (TH) | 0.009 | 0.073 | 0.017<sub>43.4</sub> | 0.898 |
| Sex (Male) | -0.011 | 0.072 | 0.021<sub>57.1</sub> | 0.886 |
| Mass at Day 12 | -0.053 | 0.035 | 2.229<sub>48.2</sub> | 0.142 |
| Laying date | -0.035 | 0.016 | 4.515<sub>43.9</sub> | **0.039** |
| **GSH:GSSG ratio** |  |  |  |  |
|  | Estimate | SE | F<sub>ddf</sub> | p |
| Treatment (TH) | 0.122 | 0.149 | 0.673<sub>48.3</sub> | 0.416 |
| Sex (Male) | 0.110 | 0.139 | 0.617<sub>57.3</sub> | 0.436 |
| Mass at Day 12 | -0.081 | 0.076 | 1.123<sub>64.2</sub> | 0.293 |

## Discussion

We tested experimentally in a wild bird species the effects of maternal thyroid hormones (THs) on nestling growth, physiology and survival by egg injection using a dosage within the natural range. We found an age-dependent effect of yolk THs on growth: TH-nestlings grew faster during the second week post-hatching compared to control nestlings. We detected no main effects of yolk TH elevation on hatching or fledging success, nor on growth or oxidative status. Likewise, the durations of embryonic and nestling periods were not affected by yolk THs.

The lack of effect on hatching success is similar to a study on great tits (Ruuskanen et al., 2016). However, two comparable studies found a positive effect of yolk THs on hatching success (rock pigeons, Hsu et al., 2017; collared flycatchers, Hsu et al., 2019). The overall low hatching success (ca. 50%) in the great tit study might have masked an effect of yolk THs on hatching. However, the study on collared flycatchers, which found a positive effect on hatchability, also had a lower overall hatching success than our study (ca. 65% vs ca. 75%, respectively). One explanation for this discrepancy may be context-dependent effects of maternal hormones on the offspring as found repeatedly for yolk androgens (e.g., Muriel et al., 2015). For example, temperature or food availability may modify the costs and benefits of elevated exposure to the maternal signal, such as induced by metabolic rate and immunity. Other egg components or hormones may also interact with yolk THs (Williams and Groothuis, 2015), such as IGF1 for the development of the skeletal system and glucocorticoids for the differentiation of gut and lung (McNabb and Darras, 2015). In addition, embryos of these two closely related species may have developed separate mechanisms to metabolise and use maternal hormones. Even within a species hormone metabolism have been found to differ between embryos (Kumar et al., 2018). Species-specific effects of yolk androgens have been documented in replica studies for yolk androgens (von Engelhardt and Groothuis, 2011). Further mechanistic and comparative studies in different contexts could help in understanding the dynamics of yolk THs during incubation.

Nestlings of TH-treated eggs grew faster than controls in the second week post-hatching. This increase in growth rate did not translate into earlier fledging nor larger body mass at fledging or higher fledging success, consistent with the results in collared flycatchers. In this species yolk THs increased early growth, but this effect disappeared at the end of the nestling phase and did not seem to confer any benefit to the TH-treated nestlings around fledging (Hsu et al., 2019). Yolk THs also increased tarsus growth in collared flycatchers, but not in pied flycatchers. In a study in great tits, yolk THs had a sex-specific effect on nestling size over the whole nestling phase, reducing size in females but increasing it in males (Ruuskanen et al., 2016). Our results show no sex-dependent effect of yolk THs on growth, with males being heavier than females throughout the nestling phase. Finally, in rock pigeons, yolk THs decreased growth in males and females (Hsu et al., 2017). Altogether, although these studies consistently show an effect of yolk THs on growth, the outcome may be context- or species-dependent, as discussed above, and a comparative approach on multiple species could help understanding these differences. Assessing the potential adaptive value of increased nestling growth would require long-term monitoring of the individuals, an experimental design hard to implement on a long-distance migrant species.

The study on rock pigeons found a sex-specific effect of yolk THs on resting metabolic rate (RMR) (Hsu et al., 2017). We expected an effect of yolk THs on oxidative stress based on these results and on the growth-enhancing effects of yolk THs (Hsu et al., 2017; Hsu et al., 2019; Ruuskanen et al., 2016). However, we observed no evidence for this hypothesis, as we found no changes in antioxidant enzyme activity (GST) or in the oxidative balance (GSH:GSSG) and no increase in oxidative damage in lipids (MDA). Thus, the increased growth in nestling pied flycatchers incurred no measurable costs in terms of oxidative stress, similarly to the earlier study on collared flycatchers that reported similar levels of oxidative stress biomarkers (Hsu et al., 2019). The absence of detrimental consequences on oxidative stress may be due to the experimental design of both studies, with an increase in yolk THs within the natural range of the species. Thus, individuals may have been able to raise their antioxidant capacities to avoid oxidative damage. In the collared flycatcher study, the oxidative status was assessed after the control nestlings had experienced a faster growth than the TH-nestlings. In our study, the effect of yolk THs on oxidative status was measured while the TH-nestlings were growing faster than the control nestlings. The absence of increased oxidative stress at day 12 is therefore good evidence that the growth-enhancing effects did not incur any oxidative costs. However, due to fieldwork constraints, there are some limits to our approach. We measured oxidative status at a single time point in one tissue, and therefore lack an overview of the variation that may have happened over the course of the whole nestling phase, also in other tissues. Villanueva et al. (2013) drew some general principles on the influence of THs on oxidative stress. For example, tissues more responsive to THs, such as liver and heart, are more susceptible to oxidative stress induced by THs. Measuring oxidative stress biomarkers in blood may not be representative of the oxidative status in other tissues, such as the liver and heart. Further studies on the relationship between yolk THs, growth and oxidative stress would be highly valuable.

In conclusion, this study provides evidence for the potential benefits of elevated prenatal thyroid hormones. We found that yolk THs increase growth, but the accelerated growth incurred no extra oxidative damage, nor affected nestling survival. The study adds to the small body of literature pinpointing to the potential, yet so far largely neglected, role of thyroid hormone-mediated maternal effects. However, reviewing the scarce literature in this field reveals inconsistencies that suggest context- or species-specific effects of yolk THs. Thus, research on maternal THs would greatly benefit from further studies with the same species in different contexts. It would also benefit from comparative studies on species with different life-histories that are likely to influence the cost-benefit trade-off induced by exposure to maternal THs.

## List of symbols and abbreviations

CO: control treatment
GSH: oxidised glutathione
GSSG: reduced glutathione
GST: Glutathione S-transferase
MDA: malonaldehyde
RMR: resting metabolic rate
T_3_: triiodothyronine
T_4_: thyroxine
THs: thyroid hormones

## Acknowledgements

We thank Florine Ceccantini for her help on the field in Turku, Anne Rokka and Arttu Heinonen for their help on yolk TH analyses. We also thank Silvia Espin for providing help on the MDA assay.

## Author contributions

TS, SR and BYH designed the study. TS and SR conducted the experiments. BYH conducted molecular sexing. SR and TS conducted the oxidative stress biomarker analyses. TS analysed the data and drafted the manuscript. All authors contributed to interpreting the data and writing the manuscript.

